# Independent acquisition of short insertions at the RIR1 site in the spike N-terminal domain of the SARS-CoV-2 BA.2 lineage

**DOI:** 10.1101/2022.04.11.487924

**Authors:** Samuele Greco, Marco Gerdol

## Abstract

Although the SARS-CoV-2 variants BA.1 and BA.2 share over 30 non-synonymous substitutions in the spike glycoprotein, they show several unique mutations that were likely acquired after the split between these two major omicron lineages. One of the most intriguing mutations associated with BA.1 is the presence of the inserted tripeptide Glu-Pro-Glu within the N-terminal domain. While the functional implications of this insertion are still unclear, several other SARS-CoV-2 lineages had previously independently acquired similarly short insertions at the very same site, named RIR1. We have previously identified this site, located approximately between codon 212 and codon 216, as a hotspot of insertions, which usually involve small nucleotide sequences including three or four codons.

Here we show that similar insertion events have independently occurred at least 13 times in early 2022 within the BA.2 lineage, being occasionally associated with significant community transmission. One of these omicron sublineages, characterized by a Ser-Gly-Arg insertion in position 212, is responsible of over 2% of all SARS-CoV-2 cases recorded in Denmark, as of early April 2022. Molecular surveillance data highlight a slow but steady growth compared with the parental BA.2 lineage in all Danish regions, suggesting that the RIR1 insertion may confer a selective advantage. We report the identification of other currently circulating BA.2 sublineages showing similar insertions, whose spread should be therefore carefully monitored in the upcoming months.

## Introduction

Since its first emergence in human during the Wuhan outbreak at the end of 2019, the genome of SARS-CoV-2 has been acquiring new mutations compared with the Wuhan-Hu-1 reference sequence at a rate close to 9×10^−4^ substitutions/site/year, based on GISAID data (Shu and McCauley, 2017). Nevertheless, in line with observations previously collected in other betacoronaviruses, such mutations do not display a uniform distribution, being disproportionately located within the the spike glycoprotein, which has a key role in regulating viral entry through the interaction with receptors expressed on host cell membranes (Boni et al., 2020; Guo et al., 2020). Most specifically, the overwhelming majority of S gene mutations target the S1 subunit, which includes the receptor-binding domain (RBD), responsible of ACE-2 binding, as well as the N-terminal domain (NTD), which contributes to host cell binding and stabilizes the spike trimer.

In light of its fundamental importance in receptor binding (Hoffmann et al., 2020; Li et al., 2003) and of its high antigenicity (Ju et al., 2020; Walls et al., 2020), the S1 subunit has been characterized by strong positive selection and by a very high rate of non-synonymous substitutions (i.e. nearly 2×10^−2^ amino acid/year) during the first two years of the pandemics. Although coronaviruses have a highly efficient proofreading activity thanks to the activity of the nsp10/nsp14 complex (Ma et al., 2015), this rate of molecular evolution far exceeds those observed in the HA1 subunit of seasonal influenza, which are usually in the range of 1-10×10−3 substitutions/amino acid/year (based on GISAID data, (Shu and McCauley, 2017)).

The evolution of SARS-CoV-2 in the human population can be briefly summarized as follows. Although all the major lineages circulating worldwide in 2020 as the result of the initial exportation from the Hubei regions were characterized by the presence of a single spike mutation (D614G) compared with the reference strain (Korber et al., 2020), for several months none of them displayed significant fitness advantage over the others. Only at the end of 2020, with relevant community immunity building up in several countries, multiple novel variants of concern (VOCs) started to emerge nearly simultaneously. The alpha, beta and gamma variants, originally described in England, South Africa and Brazil, respectively, quickly replaced all other competing variants in these regions, displaying an apparent strong selective advantage. Curiously, these three early VOCs shared a single spike mutation (N501Y), which most likely plays a key role in stabilizing the spike trimer in an open conformation, enhancing ACE2 binding (Teruel et al., 2021; Zhu et al., 2021). N501Y was paired with several other NTD mutations and, in the case of beta and gamma, with convergent RBD mutations (i.e. K417T/N and E484K) overlapping known antibody epitopes. While the rapid spread of alpha is thought to be due to its higher intrinsic transmissibility (Davies et al., 2021), the mutational patterns of beta and gamma, together with their emergence in populations with high seroprevalence and the outcomes of *in vitro* studies, strongly suggested the acquisition of significant immune escape properties (Harvey et al., 2021; Sabino et al., 2021).

The evolutionary success of these three VOCs and of several locally emerging variant of interest (VOIs) was, however, not long-lived. Indeed, in the spring of 2021, following an initial spread in the Indian subcontinent, the new VOC delta started to quickly outcompete all the other variants, to the point that alpha, beta and gamma were brought on the verge of extinction by the end of the year, with delta accounting for over 98% of all SARS-CoV-2 sequenced genomes. Although delta displayed several S1 mutations (including L452R) that might have contributed to increased immune escape (Planas et al., 2021), the most significant contribution to its higher transmissibility was likely the presence of a single mutation occurring at the S1/S2 cleavage site, P681R, which enhanced the efficiency of proteolytic cleavage (Saito et al., 2021).

On November 25^th^, 2021, WHO held a press conference announcing that the fifth VOC, named omicron, was quickly spreading in South Africa, showing a concerning pattern of reinfections (Pulliam et al., 2022). Omicron replaced delta with unprecedented speed, becoming the dominant SARS-CoV-2 lineage worldwide by early 2022. Unlike previous variants, which never showed the ability to fully escape the neutralizing activity of vaccine-elicited or convalescent sera, omicron displayed extremely significant immune evasion properties, to the point that some authors have suggested the necessity to consider this VOC as belonging to a distinct SARS-CoV-2 serotype (Simon-Loriere and Schwartz, 2022). The highly divergent antigenic properties of omicron derive from the presence of a very high number of S1 mutations, several of which have been shown to mediate ACE2 recognition and/or immune escape (Cameroni et al., 2022; Mannar et al., 2022). Curiously, many of these mutations occur at sites subjected to strong purifying selection both in previous SARS-CoV-2 variants and in bat coronaviruses, suggesting that they may cooperatively interact to mitigate slight fitness deficits associated with individual substitution, with a significant impact on the structural and biological properties of the spike trimer (Martin et al., 2022a). The highly unusual mutational patter of omicron may, to some extent, also explain the preferential use of the TMPRSS2-independent and cathepsin-mediated endosomal cell entry route (Pia and Rowland-Jones, 2022), as well as the lower fusogenity and altered cell tropism displayed by this variant (Meng et al., 2022). The extreme phylogenetic divergence between omicron and all the other previous SARS-CoV-2 variants raised important questions concerning its origins, which might be either sought in complex selective processes occurring in immunocompromised patients, or in an unidentified animal reservoir (Mallapaty, 2022).

Although omicron includes three lineages, only BA.1 and BA.2 have caused major outbreaks in human, whereas BA.3 has been responsible for less than one thousand documented cases worldwide. Although BA.1 and BA.2 share several S1 mutations, they display a remarkable sequence divergence, which is roughly comparable with the divergence observed among earlier VOCs. One of the most puzzling mutations only found in one out of the two omicron sublineages is a short insertion of three codons, which encode the tripeptide Glu-Pro-Glu at position 215, within the spike NTD of BA.1. Nearly all VOCs and VOIs carry small spike deletions compared with Wuhan-Hu-1, which may either confer enhanced antibody escape or act as permissive mutations, counterbalancing the fitness cost of otherwise deleterious RBD mutations (Meng et al., 2021). These small deletions independently occurred on multiple occasions during SARS-CoV-2 evolution at Recurrent Deletion Regions (RDRs) located within the NTD, marking one of the most significant signatures of convergent evolution documented during the first two years of the pandemics. On the other hand, the occurrence of spike insertions has been much rarer and consequently subjected to far less study. We have previously shown that insertions of short sequence stretches, usually encoding 3 or 4 amino acids, had been independently acquired over 50 times at the very same NTD site, named Recurrent Insertion Region 1 (RIR1). Before the emergence of BA.1, RIR1 insertions have been documented in alpha, gamma and delta, where they did not lead to significant community spread, and with two lineages (A.2.5 and B.1.214.2) that became dominant in a few Central America and Central African countries for a few months during 2021 (Gerdol et al., 2022). Although the functional role of RIR1 insertions is presently unclear, their convergent acquisition by multiple viral lineages most certainly suggests the need for increased monitoring, due to the possibility of associated fitness advantages.

While the frequency of observation of BA.1 is progressively declining as the result of the higher transmissibility of BA.2, we here show that a few BA.2 sublineages have independently acquired RIR1 insertions. Molecular surveillance data suggests that one of these sublineages in particular, hereby named BA.2+ins(L), which carries a Ser-Arg-Gly insertion at position 212 and is relatively widespread in Denmark, may display a moderate growth advantage compared with the parental BA.2 lineage.

## Materials and Methods

All SARS-CoV-2 genome sequence data deposited in GISAID (Shu and McCauley, 2017) up to April 11^th^, 2022, was screened, looking for sequences belonging to the BA.2 lineage and showing insertions between positions 210 and 220 in the S gene. Genomes displaying unidentified amino acids (i.e. X) in such insertions were flagged as the likely product of mis-assembly and discarded. Similarly, genomes assigned to the BA.2 lineage but carrying an “EPE” insertion in position 215, previously described as one of the lineage-defining mutations of the sister lineage BA.1, were also flagged as likely misclassified and were therefore discarded. All resulting genomes were grouped based on the inserted nucleotide sequence, whose exact position and phase of insertion were defined based on the alignment with the reference SARS-CoV-2 genome sequence (Wuhan-Hu-1, NCBI accession ID: NC_045512.2) and a reference BA.2 genome (EPI_ISL_8128502). Following the nomenclature scheme proposed in our previous work (Gerdol et al., 2022), each of these groups was labeled with progressive Roman numerals, following their chronological order of identification, starting from insertion L.

The molecular surveillance data from Denmark was subjected to further analysis, due to the local spread of one of the BA.2 sublineages carrying an insertion at RIR1, i.e. BA.2+ins(L). All genome data and associated metadata, deposited up to April 11^th^, 2022, were downloaded from GISAID. The share of BA.2 carrying insertion L was estimated, both at a national and at a regional scale (i.e. by separately taking into account Syddanmark, Sjaelland, Nordjylland, Midtjylland and Hovestaden) up to April 1^st^, 2022. The frequencies of observation of BA.2-ins(L) relative to all BA.2 sequenced genomes were reported the 7-day moving average, with 95% confidence intervals.

The phylogenetic tree was prepared as follows. All available SARS-CoV-2 genome sequence data deposited in GISAID up to April 5^th^, 2022, paired with metadata, were downloaded. Genomes with low sequencing coverage, those including stretches of undefined nucleotides (i.e. N) longer than 25 and those associated with incomplete collection date were discarded. Separate fasta files were created for viral genomes belonging to the following lineages: alpha; beta; gamma; delta; BA.1; BA.1.1; BA.2 (excluding genomes with RIR1 insertions; BA.2+ins(L) and BA.2+ins(LIII) (see **Table 1** and **Figure 1**); sequences not belonging to any VOC from the year 2020

**Table 1:**
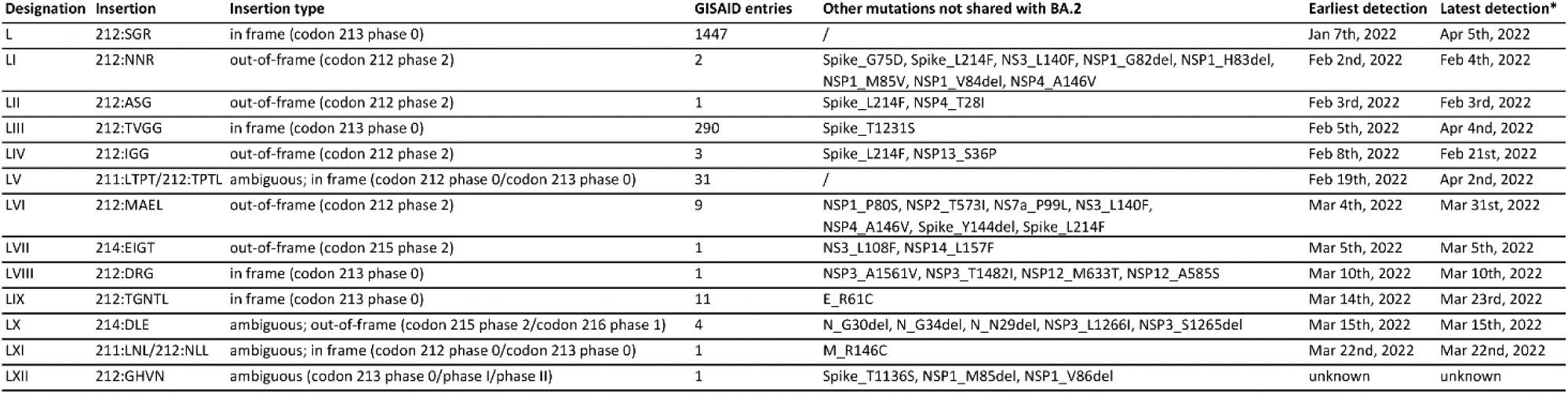
Summary of the 13 independent RIR1 insertions identified in the SARS-CoV-2 BA.2 lineage. Latest detection, as of April 11^th^, 2022.

**Figure 1:**
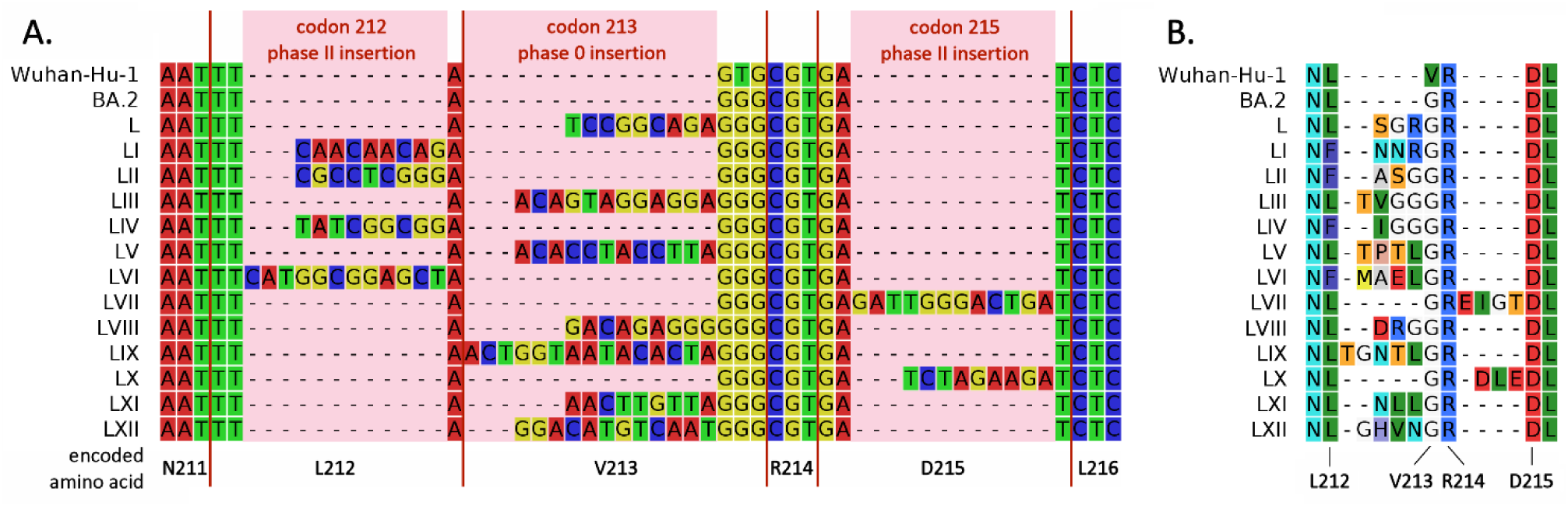
Panel A: Multiple sequence alignment of the nucleotide sequences of the S gene of viral genomes belonging to the BA.2 lineage bearing insertions at RIR1, compared with the reference sequences of Wuhan Hi-1 and BA.2. The multiple sequence alignment only shown a small region of the S gene (i.e. codon 211-codon 216). Red vertical bars highlight codon boundaries. Panel B: amino acid sequences encoded by the nucleotide sequences displayed in panel A. Codon/residue numbering refers to the Wuhan-Hu-1 reference sequence.

Particular attention was placed in an additional filtering step, which involved the fasta file including the BA.2 sequences bearing RIR1 insertions, aimed at discarding the sequences that may possibly derive from low quality assemblies. Briefly, retained sequences had to display all characterizing BA.2 mutations, defined as those present in > 80% of all sequenced genomes, as of April 5^th^, 2022. A few BA.2+ins(L) sequences displaying a small NSP1 deletion (codons 81-87) were also discarded due to its high frequency of observation in some BA.2 sublineages. Each fasta file was subsequently subsampled to include a representative group of sequences for the aforementioned lineages. In detail the final sequence dataset included: 50 alpha, beta, gamma, delta, BA.1 and BA.1.1 genomes; 150 genomes belonging to non-VOC lineages of 2020; 600 BA.2 genomes lacking RIR1 insertions; all available BA.2+ins(L) and BA.2+ins(LIII) passing quality filtering (i.e. 531 and 182, respectively). The Wuhan-Hu-1 strain genome (NCBI accession ID: NC_045512.2) was added as reference sequence

All the sampled sequences were merged and aligned with mafft v7.490 (Katoh and Standley, 2013), configured for closely related viral genomes as suggested by the authors. Gapped regions corresponding to insertions were not discarded. The best-fitting substitution model for the alignment was estimated with modeltest-ng v0.1.7 (Darriba et al., 2020). The Augur pipeline (Huddleston et al., 2021) was then used to produce the sampling date-refined phylogenetic tree, with the Wuhan-Hu-1 genome set as a reference for rooting. In order to take into account the monophyletic origin of the insertions a guide tree was used in the tree building step. All scripts and command used in this analysis are available at https://github.com/54mu/SARS-CoV-2_BA.2_RIR1insertions. The tree visualization was obtained with Auspice (Hadfield et al., 2018). Extended methods are available at https://gist.github.com/54mu/5e858e0f4aa6a7236cf5849e6ec371c7

## Results and discussion

In total, we identified 13 independent events of insertion at RIR1 associated with the BA.2 lineage. These, based on the naming scheme we have previously proposed (Gerdol et al., 2022), were chronologically ordered, from the least to the most recent one and assigned Roman numerals, from insertion L to insertion LXII (**Table 1**).

As all previously described RIR1 insertions, those associated with BA.2 involve the acquisition of either three (seven cases), four (five cases), or five (a single case) codons. However, their position was somewhat different compared with those that arose in other variants throughout 2020 and 2021 (**Figure 1A**). Indeed, four insertions (i.e. LI, LII, LIV and LX) occurred within codon 212 at phase II and four others (i.e. L, LIII, LVII and LVIII) occurred within codon 213 at phase 0. None of the 49 RIR1 insertions described in our previous work was observed in these locations (Gerdol et al., 2022). On the other hand, a single insertion (i.e. LVI) was found within codon 215, at phase II, a placement consistent with previously described RIR1 insertions. The exact location of the four remaining BA.2-associated RIR1 insertions could not be ascertained due to the possibility of multiple alternative sequence alignments with the BA.2 reference.

Due to the unusual placement of such insertions in the spike CDS, the inserted tri-, tetra- or penta-peptides in BA.2 would be usually placed between Leu212 and Val213 (albeit this residue is replaced by Gly in BA.2), and only in two cases between Arg214 and Asp215 (**Figure 1B**). Because of the out-of-frame nature of several insertions, in four cases the residue flanking the insertion at the N-terminal side (i.e. Leu212) was replaced with Phe (insertions LI, LII, LIV and LX) (**Figure 1B**). As of note, due to of the presence of the Val213Gly substitution in BA.2, the placement of most of these insertions by GISAID was often offset by a single amino acid (e.g. insertion L is reported as ins213GRG instead of ins212SGR). Consequently, a non-synonymous substitution was often incorrectly called in position 213 (e.g. Val213Ser instead of Val213Gly for insertion L). Only three out of the 49 RIR1 insertions described in our previous work were associated with a significant community spread, either locally, or globally: insertion III (lineage A.2.5), insertion IV (lineage B.1.214.2) and insertion XLI (lineage) BA.1. The first one became dominant in several Central American countries in early 2021, leading to several smaller clusters of infection abroad. The second one likely spread in central Africa in late 2020, before its importation to Belgium and Switzerland led to a few major European clusters of infection in early 2021. However, the spread of A.2.5 and B.1.214.2 was transient and neither of the two variants ever reached a global distribution. Indeed, over the following months both lineages, which predated the emergence of alpha and delta, progressively disappeared, being outcompeted by fitter variants. On the other hand BA.1, one of the two major omicron sublineages and the third major SARS-CoV-2 lineage bearing an insertion at RIR1, quickly became dominant worldwide, leading to several dozen million cases of infections and rapidly replacing delta in virtually any location.

The other 46 RIR1 insertions described in our previous work remained confined to small clusters counting no more than a few dozen sequenced cases, even though on some occasions (e.g. lineage B.1.639) undocumented community transmission likely continued for several months. Overall, these observations pointed out that the association between RIR1 insertions and the acquisition of increased fitness over previous variants could not be generalized. Due to the lack of structural overlap between RIR1 and important NTD antibody epitopes and in light of the frequent association between these insertions and non-synonymous mutations targeting the RBD, in our interpretation RIR1 insertions could be interpreted as compensatory mutations. As previously hypothesized for example for the H69/V70 deletion found in many VOCs and VOIs (Meng et al., 2021), small insertions at RIR1 may be able to mitigate slight negative fitness costs associated with the presence of non-synonymous RBD mutations affecting immune escape (Gerdol et al., 2022). From this perspective, the limited community spread of most SARS-CoV-2 variants bearing RIR1 insertions was not surprising, due to the important role played by intra-host selection in shaping spike mutational patterns, since these selective forces are not necessarily expected to lead to viral lineages with increased transmissibility. The previously documented acquisition of RIR1 insertions through *in vitro* experiments, both in response to several passages in Vero cell cultures (Shiliaev et al., 2021) and after the treatment with monoclonal antibodies (Committee for Medicinal Products for Human Use, 2021), might be consistent with this view. Hence, the acquisition of a selective advantage by of RIR1 insertions-carrying viral lineages may be conditioned by the presence or absence of other spike non-synonymous mutations associated with negative fitness costs.

In total, 11 out of the 13 BA.2-associated RIR1 insertions were only associated with a very small number of cases and were therefore unlikely to be linked with BA.2 sublineages with enhanced fitness (**Table 1**), even though it cannot be excluded that some of these, identified very recently, may further spread in the coming weeks. Nevertheless, two exceptions were immediately evident, i.e. insertion L and insertion LIII (**Table 1**). The two BA.2 sublineages carrying such insertions will be hereafter named BA.2+ins(L) and BA.2+ins(LIII). BA.2+ins(L), first detected on January 7^th^, 2022, is characterized by the presence of an in-frame TCCGGCAGA insertion between codon 212 and 213, which determines the acquisition of the tripeptide Ser-Gly-Arg in position 212 (**Figure 1**). The overwhelming majority of the BA.2 genomes deposited in GISAID carrying this insertion derives from Denmark (96.5%), the country where this sublineage apparently first originated, followed by the United Kingdom (0,9%), Germany (0,7%) and Australia (0.6%). As of April 11^th^, 2022, a few cases have been also detected in Austria, Belgium, Germany, Israel, Italy, Norway, South Africa, Sweden and Switzerland, likely as the result of exportation from Denmark. A detailed analysis of the dynamics of the spread of the BA.2+ins(L) sublineage in this country reveled an apparent slow but steady increase in frequency of observation compared with the parental BA.2 lineage over the past two months and a half (**Figure 2**). This trend is evident at a national level, where this BA.2 sublineage accounts for approximately 2.5% of all BA.2 cases as of April 1^st^, 2022 (**Figure 2A**).

**Figure 2:**
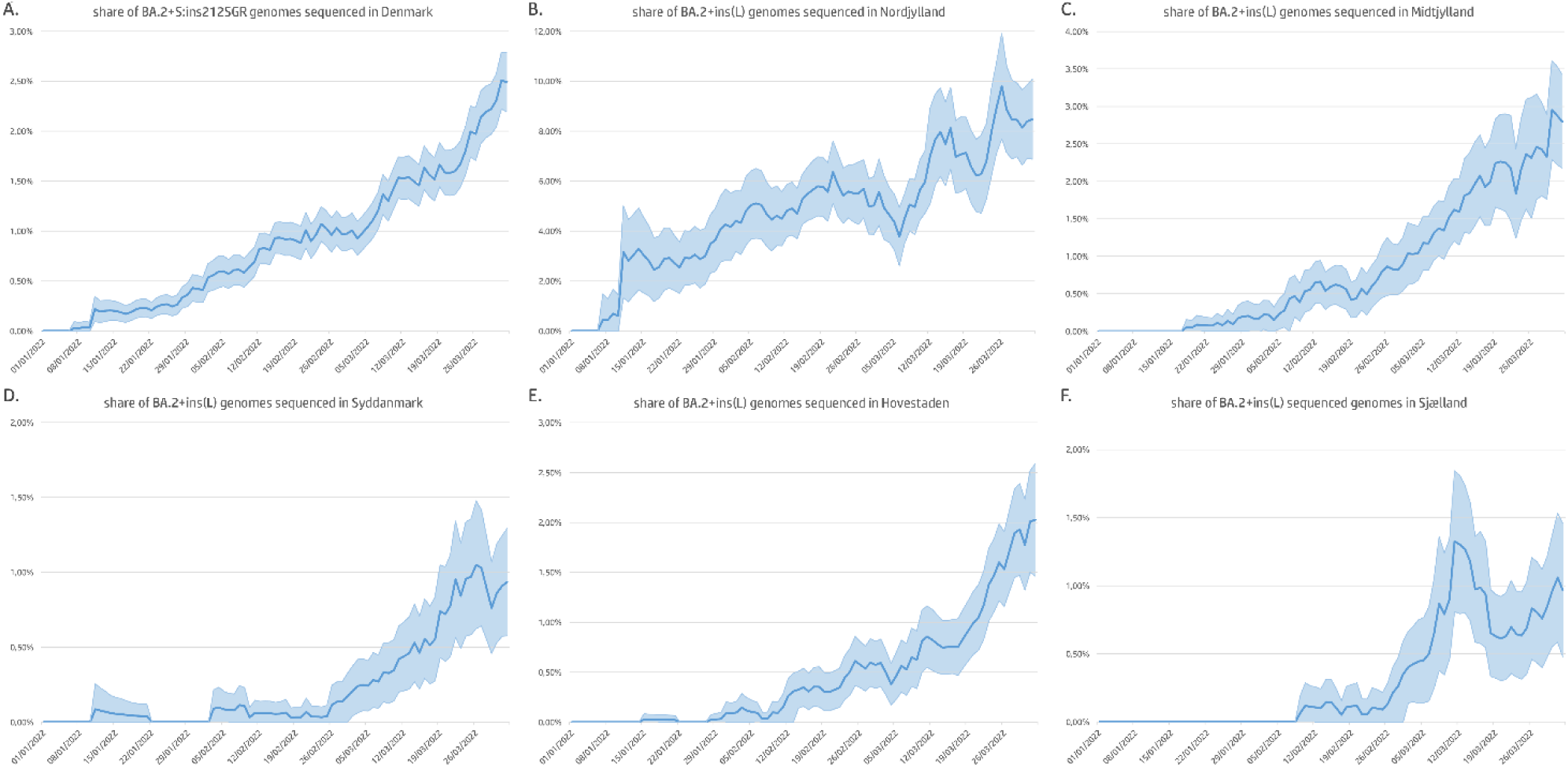
Share of BA.2 genomes bearing the L insertion at RIR1 in Denmark (panel A), with a regional break down for Nordjylland (panel B), Midtjylland (panel C), Syddanmark (panel D), Sjaelland (panel E) and Hovestaden (panel F). The graphs report the 7-day moving average frequencies of observations of BA.2+ins(L) genomes, relative to all BA.2 genomes, with 95% confidence intervals.

The national spread of BA.2+ins(L) was initially driven by the Nordjylland region, where this sublineage was first detected, undergoing a significant expansion relative to BA.2 from mid-January, up to now (**Figure 2B**). Based on the most regent genomic surveillance data, as of April 1^st^, 2022, BA.2+ins(L) may account for approximately 8.5% of all BA.2 cases recorded in the region. Midtjylland, the only Denmark region with land borders with Nordjylland, followed a similar trend: after the detection of the first cases in late-January, the share of BA.2+ins(L) genomes progressively increased over time and accounts for about 2.8% of the all BA.2 cases as of April 1^st^, 2022 (**Figure 2C**). Syddanmark, the southernmost continental region in Denmark, and Hovestaden, the easternmost and capital region, displayed nearly overlapping trends. Albeit the first few cases were detected in mid-January, the frequency of observation of this sublineage relative to BA.2 started to grow in February, and the share of BA.2 genomes carrying the RIR1 insertion L in the two regions on April 1^st^, 2022 is approximately 0.9% and 2%, respectively (**Figure 2D and 2E**). This insertion was reported for the first time in Sjælland, the largest insular region, only on February 12^th^, 2022. As of April 1^st^, 2022, BA.2+ins(L) accounts for approximately 1% of all BA.2 cases recorded in this region (**Figure 2F**).

At a national level, the comparative analysis of the incidence of infections due to BA.2 and BA.2+ins(L) allows to draw a preliminary estimate of the weekly growth advantage displayed by the latter, which would be in the range of 20%. Although the absolute number of infections linked with this sublineage presently remains relatively low (i.e. with the number of daily national cases recently dropping below 5,000, those linked with BA.2+ins(L) might be in the range of 50-100 per day), this moderate growth trend has been continuing for over two months, following consistent patterns in multiple Danish regions. It would be therefore important to clarify whether this apparent growth advantage over the parental BA.2 lineage is driven by an alteration of the structure and function of the spike protein or rather by other epidemiological factors, such as founder effects, as previously observed for other SARS-CoV-2 lineages in the past (Hodcroft et al., 2021).

The second relevant BA.2-related sublineage with an insertion at RIR1 was BA.2+ins(LIII). This sublineage, first detected on February 5^th^, 2022 in England, displays a longer insertion compared with BA-2+ins(L), i.e. ACAGTAGGAGGA, which is also found in-frame between codon 212 and 213, and encodes the tetrapeptide Thr-Val-Gly-Gly (**Figure 1**). With 290 sequenced genomes as of April 11^th^, 2022, insertion LIII is currently ranked the fifth most successful RIR1 insertion, after insertion III (A.2.5), IV (B.1.214.2), XLI (BA.1) and the aforementioned insertion L. During the first two months of its spread, BA.2+ins(LIII) has been almost exclusively detected in the UK (with 278 out of 281 cases being identified in England), with the exception of two single cases identified in Belgium and Israel, respectively. Although still linked with a very small absolute number of cases, this sublineage is currently displaying a very slight but constant growth over time and has been responsible for 0.2% of all cases reported in England in the second half of March 2022.

Curiously, neither these two insertions, nor the other 11 listed in **Table 1** were associated with other relevant S1 non-synonymous substitutions, questioning their previously hypothesized “compensatory” role in stabilizing otherwise slightly deleterious mutations with significant effect on immune escape (Gerdol et al., 2022). In detail, BA.2+ins(L) did not show any other associated non-synonymous mutations, neither in the S gene, nor in other genomic regions, leaving the Ser-Gly-Arg as the only sublineage-defining mutation present. The only other relevant mutation shared by all BA.2+ins(L) genomes was synonymous, i.e. C22791T (also located within the S gene), was not phylogenetically informative, as it was found in nearly one third of all BA.2 genomes. On the other hand, BA.2+ins(LIII) also displayed the presence of Thr1213Ser, a S2 non-synonymous conservative substitution rarely observed in SARS-CoV-2 prior to the emergence of BA.2 (only 62 cases had been recorded prior to December 2021), but occasionally found in BA.2. The only other two spike non-synonymous mutations linked with BA.2 RIR1 insertions were G75D in insertion LI, Y144del in insertion LX and T1136S in insertion LXII, even though several point mutations were found in other SARS-CoV-2 genes (**Table 1**).

In summary, most BA.2 associated RIR1 insertions appear to have not occurred in parallel with other mutations targeting the RBD, the polybasic site or known important epitopes recognized by neutralizing antibodies. The two major omicron lineages share a large fraction of the non-synonymous substitutions found in the S gene (i.e. 21), but each of the two also carry a dozen additional mutations not shared with the sister lineage. Some of the mutations shared by BA.1 and BA.2, if taken individually, would have been predicted to significantly decrease viral fitness. Nevertheless, they are thought to cooperatively interact by positive epistasis, mitigating the negative fitness costs associated with their presence and possibly altering some biological properties of the spike protein (Martin et al., 2022b). Insertion XLI (i.e. ins215EPE), which is included among the mutations found in BA.1 but not shared with BA.2, undoubtedly emerged after the split between the two lineages and its presence may therefore provide a fitness advantage in combination with any of the other mutations exclusively found in BA.1. Nevertheless, we did not observe any overlap between the presence of RIR1 insertions in BA.2 and the acquisition, by convergent evolution, of any spike mutation typical of BA.1. A different interpretation would therefore be needed to explain the emergence of RIR1 insertions in BA.2, as well as the apparent slight fitness advantage associated with BA.2+ins(L). While, to date, no conclusive explanation has been provided concerning the benefits associated with the presence of ins214EPE in BA.1, it is very likely that this small insertion, in combination with other NTD non-synonymous mutations, had a significant impact on the structure of the spike protein in BA.1. These alterations to the original primary sequence of the SARS-CoV-2 spike protein might have possibly contributed to the enhanced inter-domain and inter-subunit packing, as well as to the higher accessibility for ACE2 binding, driven by its higher predisposition towards an open configuration (Ye et al., 2022). From this perspective, the acquisition of RIR1 insertions such as L, LIII and others, might be also interpreted as a trait conferring increased stability to the BA.2 spike trimer. Hence, the independent acquisition of different short sequence stretches at RIR1 in the BA.2 lineage might simply depend on the presence of pre-existing BA.2-associated mutations and be completely independent on the acquisition of additional non-synonymous mutations. Undoubtedly, further structural studies would be required to clarify the effects of the acquisition of RIR1 insertions on a BA.2 genetic background in terms of the impact on the structural conformation and stability of spike trimers. A relevant consequence of the lack of other lineage-defining mutations in most BA.2 sublineages carrying RIR1 insertions is the inherent difficulty in their assignment through phylogenetic inference. As a matter of fact, indels are usually simply treated as missing characters and not appropriately weighted-in by most maximum likelihood approaches, including those currently used by the *agur* pipeline, i.e. IQ-TREE (Nguyen et al., 2015) and faststree (Price et al., 2010), and this might lead to significant statistical inconsistencies, as previously noted by other authors (McTavish et al., 2015; Warnow, 2012). Here we briefly report the relationships between BA.2+ins(L) and BA.2+ins(LIII) with BA.2. Despite their clear monophyletic origin, when no guide tree was provided to the maximum likelihood phylogenetic inference analysis, BA.2 genomes carrying insertion L were scattered across the BA.2 subtree (data not shown). This inconsistent placement was possibly due to a combination between the secondary acquisition, by convergent evolution, of other synonymous (or, more rarely, non-synonymous) substitutions relatively common in other BA.2 sublineages by BA.2+ins(L), or by the presence of reversions. Only when a guide tree was provided and all sequences not showing the full mutational profile of BA.2 were discarded, all BA.2+ins(L) were correctly placed in a monophyletic clade (**Figure 3**). On the other hand, this adjustment was not required to achieve a good support of the monophyly of the BA.2+ins(LIII) sublineage, which as mentioned above also carries the spike Thr1213Ser mutation. This highlights the fact that similar difficulties in future lineage assignments could arise for other SARS-CoV-2 lineages whose only distinctive mutations compared with their parental variant are either insertions or deletions.

**Figure 3:**
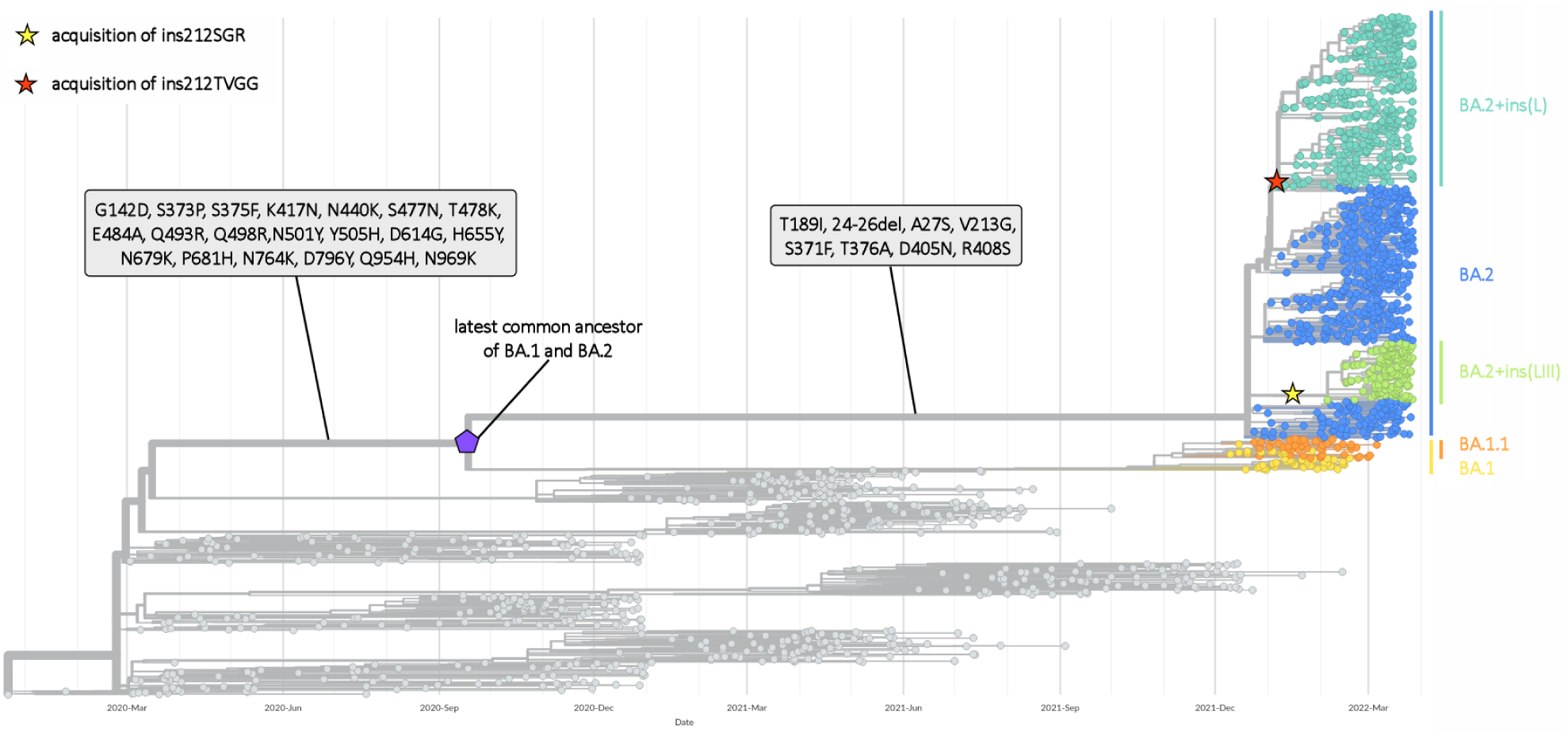
Time tree summarizing SARS-CoV-2 phylogeny and highlighting the placement of BA2+ins(L) and BA2+ins(LIII) relative with BA.1 and the other BA.2 sublineages. The acquisition of lineage-defining spike mutations in reported in grey boxes. The acquisition of insertions at RIR1 are marked by stars. The interactive tree can be visualized at https://nextstrain.org/community/54mu/SARS-CoV-2_BA.2_RIR1insertions/treeViz?c=BA_group.

## Declaration of Competing Interest

The authors declare they have no competing interests.

## Acknowledgements

The authors would like to acknowledge the tremendous efforts made by the clinicians, researchers and public health authorities that allowed the collection of SARS-CoV-2 genome data and made sequence data available in a timely manner though GISAID, as well as the great efforts made by the developers of nextstrain to assist researchers in SARS-CoV-2 evolution studies.

## Funding Statement

This research did not receive any specific grant from funding agencies in the public, commercial, or not-for-profit sectors.

## Notes

### Competing Interest Statement

The authors have declared no competing interest.

